# Unique functional neuroimaging signatures of genetic versus clinical high risk for psychosis

**DOI:** 10.1101/2024.04.03.587988

**Authors:** Charles H. Schleifer, Sarah E. Chang, Carolyn M. Amir, Kathleen P. O’Hora, Hoki Fung, Jee Won D. Kang, Leila Kushan-Wells, Eileen Daly, Fabio Di Fabio, Marianna Frascarelli, Maria Gudbrandsen, Wendy R. Kates, Declan Murphy, Jean Addington, Alan Anticevic, Kristin S. Cadenhead, Tyrone D. Cannon, Barbara A. Cornblatt, Matcheri Keshavan, Daniel H. Mathalon, Diana O. Perkins, William Stone, Elaine Walker, Scott W. Woods, Lucina Q. Uddin, Kuldeep Kumar, Gil D. Hoftman, Carrie E. Bearden

## Abstract

**Background:** 22q11.2 Deletion Syndrome (22qDel) is a copy number variant (CNV) associated with psychosis and other neurodevelopmental disorders. Adolescents at clinical high risk for psychosis (CHR) have subthreshold psychosis symptoms without known genetic risk factors. Whether common neural substrates underlie these distinct high-risk populations is unknown. We compared functional brain measures in 22qDel and CHR cohorts and mapped results to biological pathways.

**Methods:** We analyzed two large multi-site cohorts with resting-state functional MRI (rs-fMRI): 1) 22qDel (n=164, 47% female) and typically developing (TD) controls (n=134, 56% female); 2) CHR individuals (n=244, 41% female) and TD controls (n=151, 46% female) from the North American Prodrome Longitudinal Study-2. We computed global brain connectivity (GBC), local connectivity (LC), and brain signal variability (BSV) across cortical regions, testing case-control differences for 22qDel and CHR separately. Group difference maps were related to published brain maps using autocorrelation-preserving permutation.

**Results:** BSV, LC, and GBC are significantly disrupted in 22qDel compared with TD controls (False Discovery Rate q<0.05). Spatial maps of BSV and LC differences are highly correlated with each other, unlike GBC. In CHR, only LC is significantly altered versus controls, with a different spatial pattern compared to 22qDel. Group differences map onto biological gradients, with 22qDel effects strongest in regions with high predicted blood flow and metabolism.

**Conclusion:** 22qDel and CHR exhibit divergent effects on fMRI temporal variability and multi-scale functional connectivity. In 22qDel, strong and convergent disruptions in BSV and LC not seen in CHR individuals suggest distinct functional brain alterations.

## Introduction

Multiple lines of evidence suggest that disruptions to local and large-scale neural circuits are a core feature of psychosis spectrum disorders (1–3). This manifests as alterations to the temporal variability and spatial correlations of brain signals, which are detectable with resting-state functional magnetic resonance imaging (rs-fMRI) (4–9). Studying these temporal and network features of brain disruptions in people with clinical and genetic risk factors for psychosis will help us to better understand these important high-risk states and will advance our understanding of the underlying pathophysiology.

Multiple approaches can be taken to identify people at increased risk for psychosis. Schizophrenia and related disorders are highly heritable due to the polygenic effects of common genetic variants (10) as well as rarer highly penetrant mutations that include several copy number variants (CNVs) in which segments of the genome are deleted or duplicated (11). 22q11.2 Deletion Syndrome (22qDel) is one such CNV that strongly increases risk for psychosis, autism, and other neuropsychiatric disorders. This deletion of ∼46 protein-coding genes occurs in approximately 1 in 4000 people (12), of whom 10-20% can be expected to develop a psychosis spectrum disorder (13,14). Approximately 30% of 22qDel carriers endorse positive psychosis symptoms of at least moderate intensity (15–17). Highly penetrant genetic risk factors like 22qDel represent important opportunities for characterizing biological pathways underlying psychosis risk in a “genetics-first” approach (18,19). Alternatively, in a “behavior-first” framework, a Clinical High Risk (CHR) state can be defined based on the presence of sub-threshold positive symptoms, without considering specific genetic risk factors (20). Approximately 20-30% of people meeting CHR criteria can be expected to convert to a diagnosable form of psychotic disorder within a three-year follow-up period (20,21).

Substantial research has been done on various functional neuroimaging measures in separate studies of 22qDel and CHR relative to typically developing (TD) controls, with a range of convergent and divergent outcomes (22,23). However, no study to date has directly compared rs-fMRI correlates of these genetic and clinical high-risk states using the same analytic pipelines. Here, we conduct a mega-analysis of rs-fMRI data from a total of 687 22qDel carriers, CHR individuals, and age-sex- and site-matched TD controls collected at multiple sites and subjected to a harmonized preprocessing and analysis pipeline.

Our aims are to: *i)* characterize shared and unique effects of 22qDel and CHR on rs-fMRI measures of global brain connectivity (GBC), local connectivity (LC), and brain signal temporal variability (BSV), and *ii)* gain deeper biological understanding of fMRI phenotypes by relating these findings to other multimodal maps of regional brain features. The three rs-fMRI measures index different spatial and temporal aspects of neurophysiology. GBC reflects connectivity of large scale functional networks which have been shown to be disrupted in 22qDel, schizophrenia, and CHR (24–26). LC is often measured as the similarity in signal between spatially adjacent voxels, to which alterations have been observed in schizophrenia and in first degree relatives at genetic high risk (6,27–29). BSV is a heritable brain phenotype that relates to hemodynamic physiology, excitation/inhibition balance, and the spatial expression pattern of schizophrenia risk genes (30,31). We chose these three measures to capture functional brain alterations at multiple scales across the same sets of cortical regions. To generate biologically-informed hypotheses from our rs-fMRI findings, we relate spatial maps of case-control group differences to published brain maps from multiple sources, such as metabolic data from positron emission tomography (PET) and gene expression data from the Allen Human Brain Atlas (32–35).

## Methods

### Participants

The neuroimaging dataset contains a total of 687 participants from two separate multi-site studies (**Table 1**): 164 carriers of molecularly confirmed 22qDel along with 134 matched TD controls (Control-22q), and 240 individuals with CHR for psychosis plus 149 matched TD controls (Control-CHR). 22qDel and Control-22q data were shared from five scanners/sites in the United States, Europe, and the United Kingdom. CHR and Control-CHR data came from the North American Prodrome Longitudinal Study 2 (NAPLS2) (36,37), which includes eight sites in the United States and Canada. CHR participants were adolescents and young adults ages 12-35 with subthreshold psychosis symptoms (38). See **Supplemental Methods** for inclusion/exclusion criteria and a detailed description of participant counts by site. After study procedures were fully explained, adult participants provided written consent, while participants younger than 18 years provided written assent with the written consent of their parent or guardian. The respective Institutional Review Boards at each site approved all study procedures and informed consent documents.

**Table 1.**
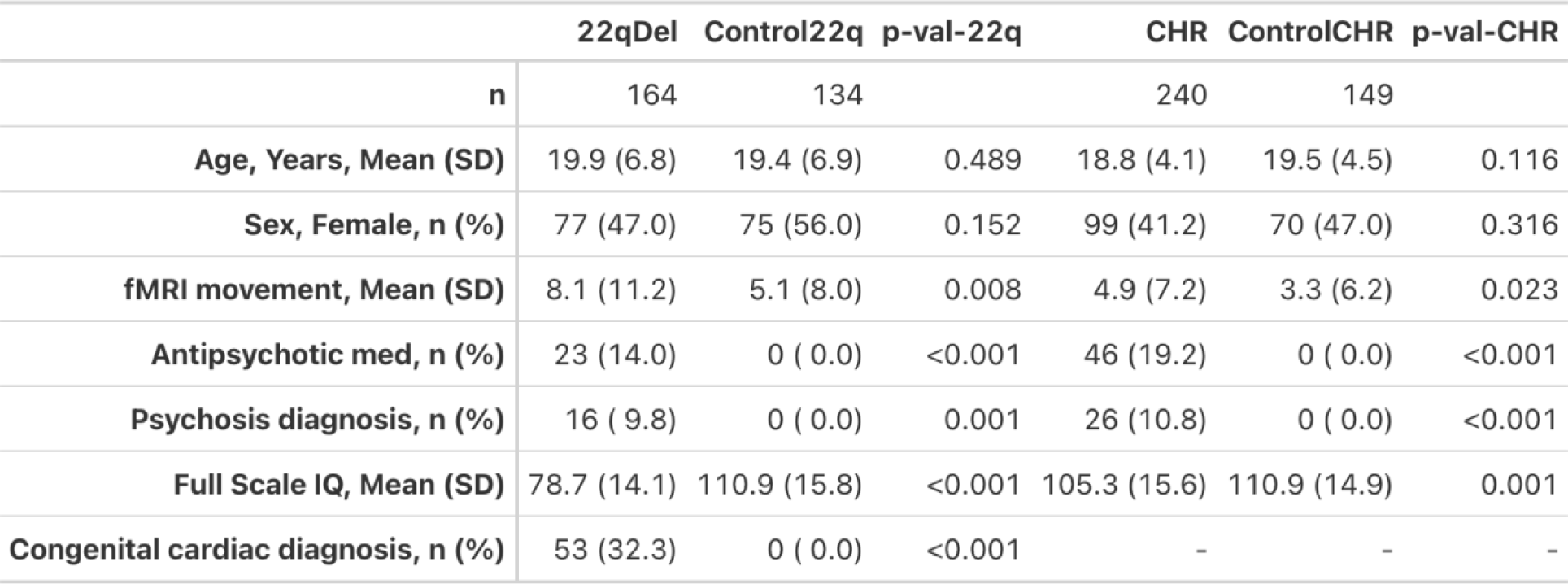
Demographics. 22q11.2 Deletion Syndrome (22qDel) carriers and matched typically developing controls (Control22q), as well as Clinical High Risk (CHR) patients and matched typically developing controls (ControlCHR). p-val-22q describes difference tests (ANOVA or chi-squared) between 22qDel and Control22q groups. p-val-CHR refers to comparisons between CHR and ControlCHR groups. fMRI movement describes the percentage of frames removed per scan for exceeding displacement/intensity thresholds. Antipsychotic med indicates the proportion of participants taking medication in that drug class. In the 22qDel group, “Psychosis diagnosis” indicates diagnosis of psychotic disorder (schizophrenia, schizoaffective, or not otherwise specified) at the time of scan. In the CHR group, “Psychosis diagnosis” indicates those who were diagnosed with a psychotic disorder at any point during the two-year study follow-up period (i.e. CHR individuals who eventually “convert” to a psychosis spectrum diagnosis). Psychotic disorder diagnosis was based on Structured Clinical Interview for DSM/SCID-5. Full Scale IQ includes normed scores from multiple scales: Wechsler Adult Intelligence Scale (WAIS), Wechsler Adult Intelligence Scale-Revised (WAIS-R), Standard Progressive Matrices (SPM), Wechsler Intelligence Scale for Children-III (WISC-III), Wechsler Abbreviated Scale of Intelligence-II (WASI-II). Congenital cardiac diagnosis indicates any 22qDel carrier with a documented diagnosis of a congenital heart defect (e.g., septal defect, Tetralogy of Fallot, etc.).

### Neuroimaging acquisition and processing

Blood oxygen level dependent (BOLD) resting-state functional magnetic resonance imaging (rs-fMRI) and high-resolution structural images were collected at each study site (see **Supplemental Methods**). All data from 22qDel, CHR, and TD controls were processed with the same workflow, as described in detail in previous publications (39,40). Functional and structural images were processed with the Quantitative Neuroimaging Environment and Toolbox (41), applying a modified version of the methods developed for the Human Connectome Project (HCP) (42), as well as motion scrubbing, i.e. censoring frames with displacement or intensity change thresholds exceeding those recommended by Power et al. (43–45). Functional connectivity analyses were computed on the residual of the signal after regression of motion time series, the mean signal time series from the ventricles and deep white matter, and the first derivatives of these measures.

### fMRI measures

Three rs-fMRI measures were calculated for each scan: global brain connectivity (GBC), local connectivity (LC), and brain signal variability (BSV). All three measures used the same set of 360 cortical regions defined from multi-modal MRI in 210 healthy young adults from the HCP (46). See **Supplemental Methods** for details on each measure. Computations were performed in R using ciftiTools to manipulate neuroimaging data (47).

GBC is a well-validated measure defined as the average functional connectivity between a given brain region and all other regions (48,49). A high GBC value indicates a region in which signal is similar to many other regions of the brain, whereas a low GBC value represents a region that is dissimilar to the majority of other regions.

LC was calculated as a measure of the cohesiveness or homogeneity of vertex-level BOLD time series within the same set of regions. This approach is based on the network homogeneity method, wherein FC is computed between each pair of voxels in a chosen network (50).

BSV was calculated as the average temporal standard deviation of the BOLD time series in each region. This measure of variability is also referred to as resting state fluctuation amplitude, and represents a heritable brain phenotype that relates to underlying excitation/inhibition balance and hemodynamic physiology (30,31).

To correct for variability related to site/scanner, we used a neuroimaging-optimized implementation of ComBat (51) which uses empirical Bayes methods to correct for batch/site effects, with increased robustness compared to linear model approaches (52). After ComBat, values for each measure were normalized within each region based on the mean and standard deviation for the relevant control group.

### Group-level fMRI comparisons

For each fMRI measure, across each region, linear models were used to test the main effect of 22qDel versus matched controls (Control22q), and the main effect of CHR versus matched controls (ControlCHR). 22qDel and CHR groups were compared to their respective control groups but not directly to each other to avoid the confounding effects of scanner/site and demographic differences between the 22qDel and CHR groups. All models controlled for linear and quadratic age, sex, site, and movement (measured as the percentage of frames scrubbed from each scan), and p-values for the main effect of group were adjusted for multiple comparisons with False Discovery Rate (FDR) *q*<0.05 for each brain map (53). See **Supplemental Methods** for more details on linear models.

### Brain map comparisons

In order to assess similarity of the three fMRI measures within and between clinical groups, we used permutation methods to compare the spatial brain maps for case-control comparisons with significant group main effects. For a given pair of maps, similarity was tested via Pearson’s correlations, and two-tailed p-values were computed from a null distribution generated from 10,000 spatial autocorrelation-preserving surrogate brain maps per hemisphere, generated with BrainSMASH (33), see **Supplemental Methods**.

We next used the same permutation testing procedure to compare the left hemisphere cortical maps from our case-control analyses to a set of 22 left hemisphere cortical maps from previously published datasets, using the neuromaps and abagen toolboxes (35,54), see **Supplemental Methods**. Maps include metabolic and physiologic data from positron emission tomography (PET), gene expression from post-mortem tissue, and various measures from magnetoencephalography (MEG), structural MRI, and functional MRI; see **Table 2** for a description of each map (34,46,55–61). Analyses were restricted to the left hemisphere because some datasets did not include densely sampled right hemisphere data.

**Table 2.** Reference brain maps. Description of neuroimaging modality and citation for each published brain map (34,46,55–61). PET = positron emission tomography; mRNA = messenger ribonucleic acid; fMRI = functional magnetic resonance imaging; sMRI = structural MRI; MEG = magnetoencephalography; PC1 = first principal component; PVALB-SST = difference in normalized parvalbumin and somatostatin expression; PNC = Philadelphia Neurodevelopmental Cohort; NIH = National Institute of Health; T1w = T1-weighted sMRI; T2w = T2-weighted sMRI.

| Map | Modality | Citation |
| --- | --- | --- |
| Glucose metabolism | PET | Vaishnavi et al. 2010 |
| Oxygen metabolism | PET | Vaishnavi et al. 2010 |
| Cerebral blood flow | PET | Vaishnavi et al. 2010 |
| Cerebral blood volume | PET | Vaishnavi et al. 2010 |
| PC1 gene expression | mRNA | Markello et al. 2021 |
| PVALB-SST gradient | mRNA | Markello et al. 2021 |
| fMRI gradient | fMRI | Margulies et al. 2016 |
| Intersubject variability | fMRI | Mueller et al. 2013 |
| PC1 neurosynth | fMRI | Yarkoni et al. 2011 |
| Allometric scaling (PNC) | sMRI | Reardon et al. 2018 |
| Allometric scaling (NIH) | sMRI | Reardon et al. 2018 |
| Developmental expansion | sMRI | Hill et al. 2010 |
| Evolutionary expansion | sMRI | Hill et al. 2010 |
| Cortical thickness | sMRI | Glasser et al. 2016 |
| T1w/T2w ratio | sMRI | Glasser et al. 2016 |
| MEG delta | MEG | Van Essen et al. 2013 |
| MEG theta | MEG | Van Essen et al. 2013 |
| MEG alpha | MEG | Van Essen et al. 2013 |
| MEG beta | MEG | Van Essen et al. 2013 |
| MEG gamma1 | MEG | Van Essen et al. 2013 |
| MEG gamma2 | MEG | Van Essen et al. 2013 |
| ROI size | sMRI | Glasser et al. 2016 |

### Clinical and cognitive analyses

The primary CHR fMRI analyses were repeated comparing the 26 CHR individuals who eventually converted to a psychosis diagnosis (CHRc) to the ControlCHR group, and to the subset of the CHR participants who did not convert. All models controlled for age, age^2^, sex, site, and movement.

The primary 22qDel fMRI analyses were repeated comparing the 54 22qDel carriers with psychosis risk symptoms (PS+) and the 110 22qDel carriers without (PS-), controlling for age, age^2^, sex, site, and movement. See **Supplemental Methods** for classification of psychosis risk status based on clinical scales.

In an exploratory analysis of cognition, linear relationships were tested for each measure in each group across all regions between fMRI measures and Full Scale IQ controlling for sex, site, and movement.

For all clinical and cognitive analyses, significance for the relevant main effects were evaluated at FDR q<0.05 within each set of tests for a given group and measure.

### Secondary analyses

For all regions with a significant main effect of group in the original model, p-values for the main effects of site were investigated.

The primary case-control analyses did not include global signal regression (GSR) as a preprocessing step for consistency with prior NAPLS work (26,62) and due to systemic associations between global signal topography and factors such as age (63). However, case-control comparisons were repeated with the additional inclusion GSR. Within the 22qDel and CHR groups, across all regions, linear models were tested for the relationship between fMRI measures and antipsychotic medication status.

Because congenital heart disorders are common in 22qDel (**Table 1**), we tested a linear model for fMRI effects of any congenital cardiac diagnosis (e.g. atrial or ventricular septal defect, conotruncal defect, etc.) within the 22qDel group.

For all secondary analyses, significance for the relevant main effects were evaluated at FDR q<0.05 within each set of tests for a given group and measure. All models controlled for age, age^2^, sex, site, and movement.

## Results

### fMRI analyses

22qDel carriers significantly differ from matched controls with respect to each fMRI measure (GBC, LC, and BSV); **Figure 1a-c**. GBC is lower in 22qDel for a set of brain regions including bilateral somatomotor, visual, and temporal cortex. No areas of higher GBC in 22qDel survived correction for multiple comparisons. Both LC and BSV show decreases in 22qDel relative to controls across a similar set of frontal and parietal regions, and similar increases in inferior temporal regions. However, LC is decreased in somatomotor cortex, which is not observed for BSV in 22qDel.

**Figure 1.**
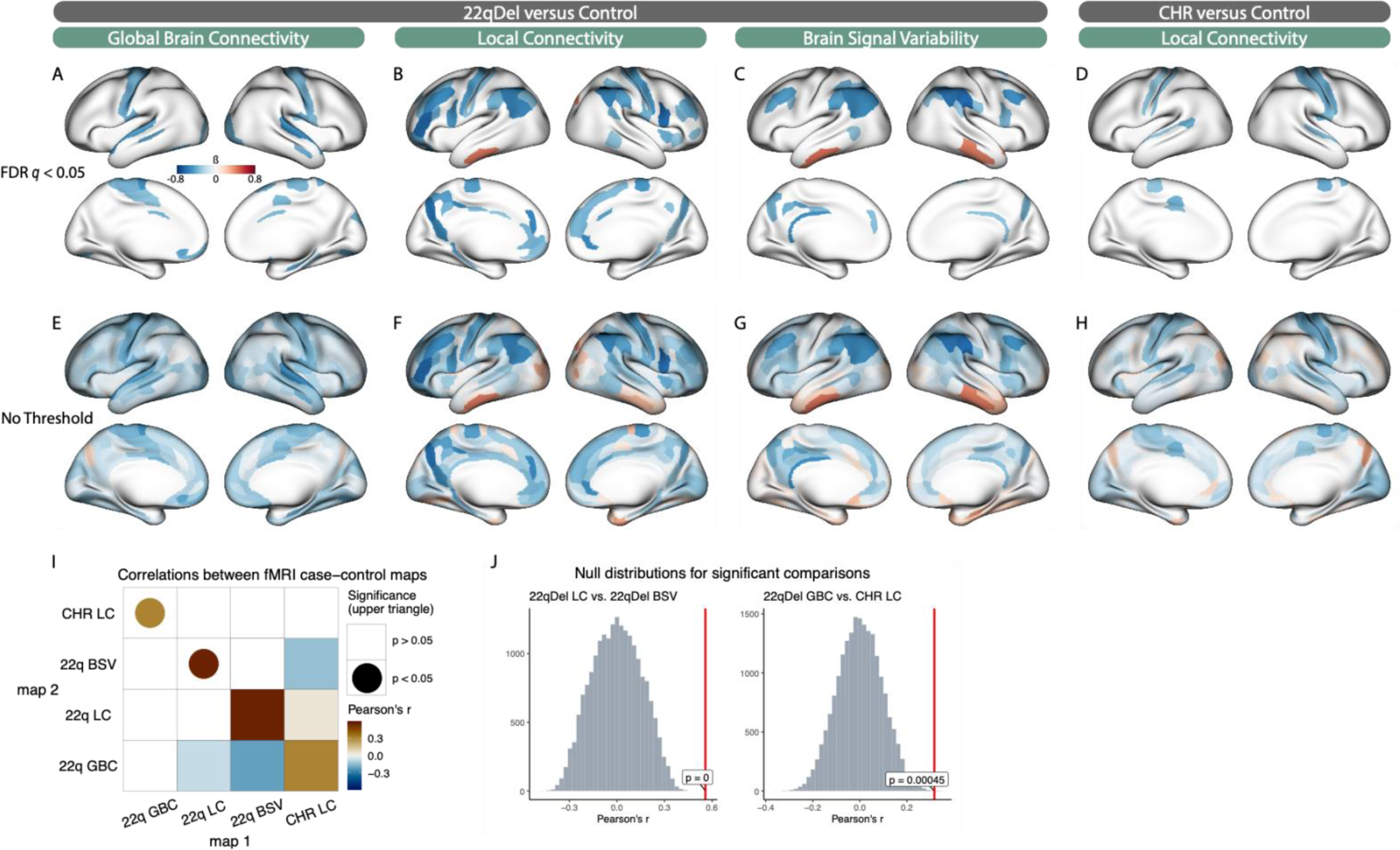
Case-control fMRI differences. **A-C**) Standardized coefficients for significant main effects of 22qDel versus controls for global brain connectivity (GBC), local connectivity (LC), and brain signal variability (BSV), visualizing effect sizes for regions where False Discovery Rate (FDR) corrected *q* < 0.05. Blue indicates 22qDel < control, red indicates 22qDel > control. **D**) regions with significantly decreased LC in clinical high risk (CHR) versus controls. No significant differences in GBC or BSV were found between CHR and controls. **E-H**) Same as **A-D** visualizing standardized group difference effect sizes for all cortical regions (no significance threshold). **I**) Correlations between brain maps in **E-H**. All correlations are shown in the lower triangle; upper triangle shows only those with p<0.05. **J**) Two-tailed p-values are calculated from the true correlation between brain maps (red line) compared to a null distribution (gray histogram) of correlations to 10,000 spatial autocorrelation-preserving surrogate brain maps per hemisphere.

In CHR individuals compared to matched controls, there are no significant differences in GBC or BSV. However, local connectivity is significantly decreased in CHR relative to controls in a set of somatomotor and temporal regions (**Figure 1d**).

Testing correlations between the four threshold-free brain maps with significant group effects (22qDel GBC, LC, and BSV, and CHR LC; **Figure 1i**), using spatial autocorrelation-preserving permutations, revealed that the effect of 22qDel is similar between LC and BSV (driven by similarity in frontal-parietal and temporal regions; **Figure 1b,c**), and that the effect of 22qDel on GBC resembles the effect of CHR on LC (driven by somatomotor regions; **Figure 1a,d**). While the threshold-free maps for group effects on LC are not significantly correlated between 22qDel and CHR, there is a notable overlap of significant effects, in which 6 somatomotor regions show significant effects for both CHR and 22qDel (out of 16 total significant regions in CHR).

### Multi-modal brain map relationships

The map of 22qDel versus control differences for BSV is significantly related to published PET maps of glucose and oxygen metabolism as well as cerebral blood flow and volume (55), such that decreased BSV in 22qDel is associated with regions of typically high metabolism and blood flow. 22qDel BSV is also significantly correlated with a map of inter-individual variability from an rs-fMRI study of typical adults (57). The map of LC differences in 22qDel versus controls is similarly related to cerebral blood flow and volume (55). LC in 22qDel was also related to the principal sensory/associative gradient extracted from a large rs-fMRI study, highlighting default mode and frontoparietal regions with low LC in 22qDel (56). The 22qDel LC map is also related to region of interest (ROI) size in the atlas. All relationships had the same direction of effect, in which lower values in the 22qDel effect size map (representing 22qDel < control) are related to higher values in the reference map; see **Figure 2**. See **Table 2** for descriptions of each reference map. Brain map comparisons for the CHR group are shown in **Supplementary Figure S1**. Similar to 22qDel, ROI size was significantly related to the CHR versus ControlCHR difference map for local connectivity. No other LC brain map relationships were found to be significant in the CHR analysis. The effects of CHR on GBC and BSV were not strong enough to allow interpretable brain map comparisons.

**Figure 2.**
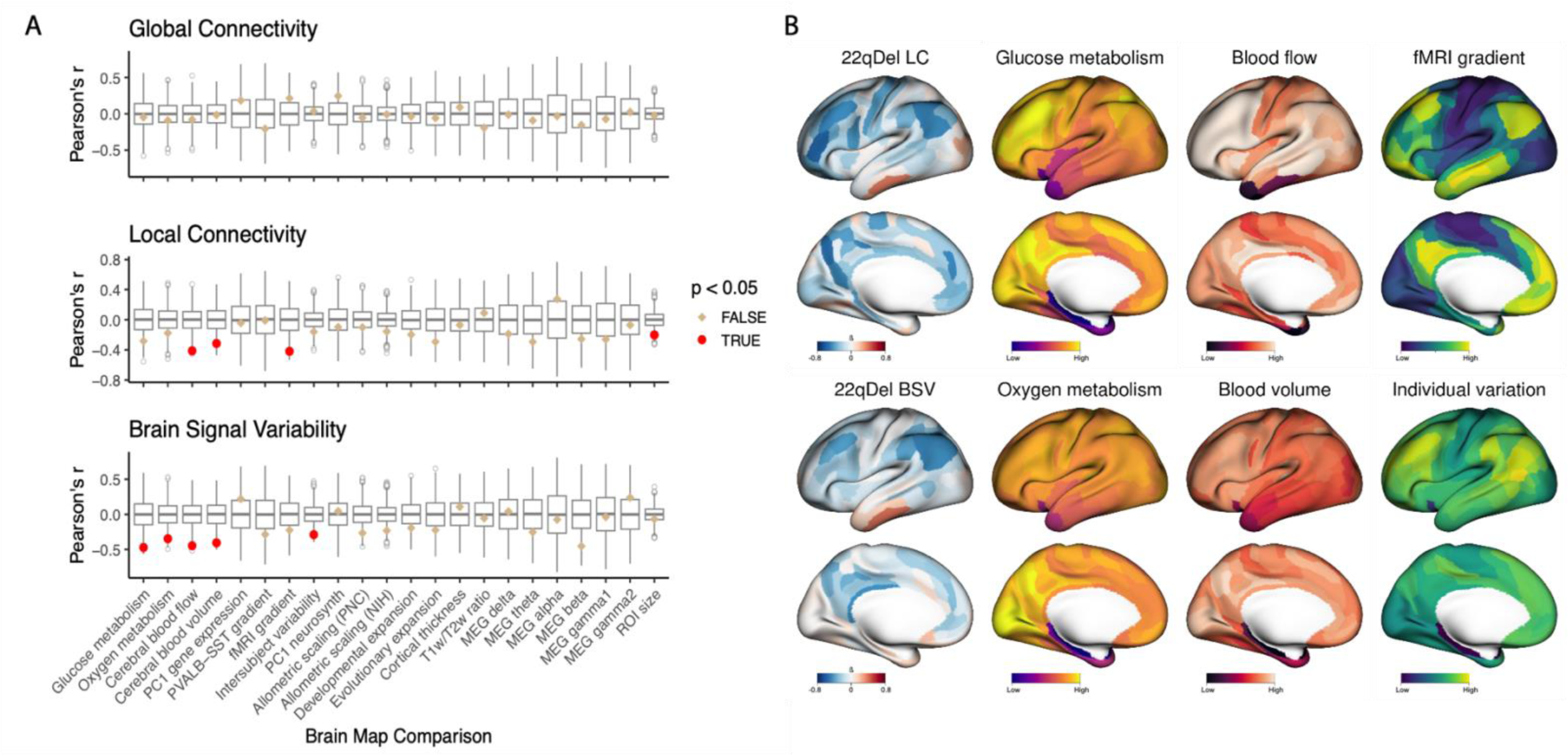
Multi-modal brain map comparisons for 22qDel fMRI effects. **A**) Left hemisphere cortical maps from 22qDel case-control models were tested for spatial similarity to multiple publicly available datasets including metabolic and physiologic data from positron emission tomography (PET), gene expression from post-mortem tissue, and various measures from magnetoencephalography (MEG), structural magnetic resonance imaging (MRI), and functional MRI. Null distributions (gray box plots) were computed from the Pearson correlations between 10,000 spatial autocorrelation-preserving permutations of the target fMRI 22qDel-control map (named in the plot title) and the second map of interest (named on the x-axis). The true correlation values between the two maps of interest are marked by tan or red points, and two-tailed p-values are computed from the proportion of values in the null distribution whose absolute value exceeds the absolute value of the true test statistic. **B**) Left hemisphere maps for 22qDel-control group differences in local connectivity (LC) and brain signal variability (BSV), as well as example maps for glucose and oxygen metabolism (55), cerebral blood flow and volume (55), and fMRI maps describing a sensory/transmodal gradient (56) and inter-individual variation (57).

### Clinical and cognitive analyses

Across all three fMRI measures, there were no regions where we observed a significant main effect of CHRc versus ControlCHR, or CHRc versus CHR. Similarly, in 22qDel there were no regions with a significant effect of PS+ versus PS-.

In an exploratory analysis of cognition, there were no regions with significant linear relationships between Full Scale IQ and any fMRI measure in any group (22qDel, Control22q, CHR, ControlCHR).

### Secondary analyses

For all regions with a significant main effect of group, none showed a significant effect of dummy coded site for any measure in the 22qDel and CHR analyses. In fact, the uncorrected p-values for site effects in these regions were all greater than 0.05. This provides reassurance that after ComBat correction site/scanner differences are not driving the observed results.

Within the 22qDel and CHR groups, there were no brain regions with significant main effects of antipsychotic medication status in any of the three measures. This suggests that our measures are more sensitive to case-control differences than medication effects.

Within the 22qDel group, across all three rs-fMRI measures, there were no regions with a significant main effect of congenital heart defect diagnosis.

Across both clinical groups and all three measures, the results of case-control analyses with the inclusion of GSR are highly consistent with our initial findings that did not include GSR (see **Supplemental Figure S3**).

## Discussion

This is the first study to compare patterns of functional brain disruption in the clinical high-risk (CHR) and genetic high-risk (22qDel) conditions. We find robust and significant effects of 22qDel versus matched controls across all three measures: global brain connectivity (GBC), local connectivity (LC), and brain signal variability (BSV) (Figure 1). The CHR group only differs significantly from controls for LC. Overall, the rs-fMRI effects of 22qDel and CHR are mostly dissimilar. Comparison to previously established brain maps suggests that spatial patterns of 22qDel versus control differences in LC and BSV relate to multiple patterns including regional variation in cerebral blood flow and metabolism, measured from PET studies in neurotypical individuals.

### Case-control findings

22qDel and CHR both show significantly decreased LC across a set of somatomotor regions, but the overall spatial patterns of LC differences from controls are not significantly correlated between the two groups. Somatomotor LC has not been directly characterized before in 22qDel or CHR, as we did here; however, using other analytic approaches functional connectivity between somatomotor cortex and the thalamus has been found to be increased in both 22qDel youth and in CHR individuals who convert to a psychotic disorder (39,40,64).

In 22qDel, LC and BSV can be seen to converge on a similar pattern of disruptions: a set of frontal and parietal regions show both decreased temporal variability and decreased within-region spatial homogeneity (i.e., LC), with the opposite pattern observed for inferior temporal regions.

We do not find any significant differences between the CHR and control groups in terms of BSV. Prior studies in schizophrenia have found evidence of widespread increases in fMRI signal variance relative to controls (4,5). In CHR, the fractional amplitude of low frequency fluctuations has been shown to differ from controls (65), suggesting that frequency-specific measures of signal variability may be more sensitive to the CHR phenotype.

Similarly, we do not find significant CHR effects on GBC. Graph theory based approaches have found functional connectivity differences between CHR and control individuals (26,66), which our GBC measure does not capture. In this CHR sample we also do not see the GBC differences in prefrontal cortex and other regions that have been observed in schizophrenia (4,49,67). This suggests that functional connectivity disruptions on smaller spatial scales, or between specific long-range nodes, may be more indicative of the CHR state, but that overall global connectivity of regions is not as strongly impacted.

### Multi-modal brain map relationships

For brain-wide case-control fMRI effects, we evaluated relationships to multiple published brain maps (**Table 2, Figure 2, Supplemental Figure S2**). Regions with significantly decreased LC and BSV in 22qDel relative to controls are both associated with high cerebral blood flow and blood volume from a PET study of typical adults (55). The map of BSV differences in 22qDel was additionally associated with glucose and oxygen metabolism from the same PET study. While these are exploratory findings, they are consistent with the hypothesis that neuronal metabolic pathology and/or vascular and hemodynamic abnormalities in 22qDel underlie some of the observed rs-fMRI differences. 22qDel is associated with high rates of congenital cardiac defects (12) and increased neurovascular anomalies (68,69). In our sample, we did not find a relationship between congenital heart defects and rs-fMRI measures, but there may be neurovascular alterations in 22qDel that are unrelated to categorical cardiac defect status. A recent arterial spin labeling MRI study showed increased cerebral blood flow in 22qDel compared to controls (70). Additionally, animal models of 22qDel and studies of induced pluripotent stem cell-derived neurons from human 22qDel samples provide convergent evidence for neuronal mitochondrial disruption (71–74) and disrupted neurovascular development (75–78) caused by 22q11.2 CNVs. Neurovascular interactions are fundamental to the BOLD fMRI signal (79,80), and play a key role in neurodevelopmental processes such as cortical expansion (76,81,82), which appears to be strongly impacted in 22qDel given structural MRI findings of markedly reduced cortical surface area (83).

22qDel LC effects were also related to the principal gradient of functional connectivity from a study in typical adults (56), and BSV effects were related to intersubject variability in another study of typical adults (57). This is a reflection of the preferential LC and BSV decreases in 22qDel across a distributed network of highly dynamic association regions in frontal, parietal, and temporal cortex. The disrupted local connectivity and variability of the default mode and related associative networks converge with prior findings of decreased functional and structural connectivity in the default mode network in 22qDel (24,84).

For both 22qDel and CHR, LC was also found to be preferentially decreased in larger regions (e.g. somatomotor regions). No other significant brain map relationships were found for CHR effects on LC. No significant relationships were found for effects of 22qDel or CHR on GBC, and results should not be interpreted for the effect of CHR on BSV because the effect sizes are too small to facilitate comparison (**Supplemental Figures S1-S2**).

### Strengths, limitations, and future directions

A significant strength of this work is that preprocessing and analytic workflows have been harmonized across all data from 22qDel, CHR, and control groups. Our mega-analytic approach leverages data from multiple sites to increase statistical power and generalizability, while minimizing methodological differences as a source of non-biological variation which can obscure true signals. This allows us to compare 22qDel and CHR findings with greater clarity than ever before. Our secondary analyses provide evidence that our findings are not strongly influenced by site/scanner differences or antipsychotic medication status in either group. Encouragingly, we also find that all case-control rs-fMRI results are highly comparable with and without the inclusion of GSR as an additional denoising step.

Despite our relatively large sample size, we still have limited power to detect effects specific to the subsets of the 22qDel and CHR samples with a psychosis diagnosis. Only 26 of the 240 CHR participants converted to a psychosis diagnosis during follow-up, and only 16 of the 164 22qDel participants in our young cohort had a diagnosis of a psychotic disorder. A larger proportion of the 22qDel group met criteria for subthreshold psychosis risk symptoms (54/164), but this constitutes a heterogeneous group, and they did not show rs-fMRI differences from the 22qDel carriers without psychosis symptoms. While the robust rs-fMRI effects we see are reflective of the CNV broadly, they may not differentiate people with varying psychiatric phenotypes. However, future studies with larger samples may uncover psychosis-specific rs-fMRI effects. We also did not find any significant relationships between Full Scale IQ and rs-fMRI, but future large studies with standardized and detailed cognitive data will be better suited to analyses of brain-behavior relationships.

The brain map comparison analyses are exploratory and intended to generate rather than confirm hypotheses. Our findings of strong relationships between 22qDel-specific disruptions and gradients of brain hemodynamic and metabolic activity warrant further research into the underlying neurophysiology. fMRI studies with additional physiological measurements such as breathing belts or capnography, or tasks such as breath-holding studies in 22qDel could help disentangle fMRI correlates of altered neuronal activity from those relating to altered hemodynamics and blood oxygenation.

### Conclusions

We show, for the first time, that 22qDel carriers and individuals at CHR for psychosis exhibit highly distinct alterations in rs-fMRI measures of global brain connectivity, local connectivity, and temporal variability. Compared to controls, 22qDel carriers show marked disruptions across all three measures, whereas CHR individuals differ only in local connectivity. Comparison to multi-modal brain maps suggests that, uniquely in 22qDel, temporal variability and spatial homogeneity of rs-fMRI signals are preferentially reduced in association cortex regions with high hemodynamic and metabolic activity. Findings motivate future research to characterize points of convergence between CHR and genetic risk syndromes, as well as specific research into the neurovascular and neurometabolic underpinnings of functional brain alterations in 22qDel.

## Acknowledgments

**Charles H. Schleifer:** Conceptualization, Methodology, Software, Formal Analysis, Data Curation, Writing - Original Draft, Writing - Review & Editing, Visualization

**Sarah Chang:** Data Curation, Methodology, Writing - Review & Editing

**Carolyn Amir:** Data Curation, Methodology, Writing - Review & Editing

**Kathleen P. O’Hora:** Data Curation, Methodology, Writing - Review & Editing

**Hoki Fung:** Data Curation, Methodology, Writing - Review & Editing

**Jee Won D. Kang:** Data Curation, Methodology, Writing - Review & Editing

**Leila Kushan-Wells:** Data Curation, Project Administration

**Eileen Daly:** Data Curation, Project Administration

**Fabio Di Fabio:** Data Curation, Project Administration

**Marianna Frascarelli:** Data Curation, Project Administration

**Maria Gudbrandsen:** Data Curation, Project Administration

**Wendy R. Kates:** Data Curation, Project Administration

**Declan Murphy:** Data Curation, Project Administration

**Jean Addington:** Data Curation, Project Administration

**Alan Anticevic:** Data Curation, Project Administration, Software

**Kristin S. Cadenhead:** Data Curation, Project Administration

**Tyrone D. Cannon:** Data Curation, Project Administration

**Barbara A. Cornblatt:** Data Curation, Project Administration

**Matcheri Keshavan:** Data Curation, Project Administration

**Daniel H. Mathalon:** Data Curation, Project Administration

**Diana O. Perkins:** Data Curation, Project Administration

**William Stone:** Data Curation, Project Administration

**Elaine Walker:** Data Curation, Project Administration

**Scott W. Woods:** Data Curation, Project Administration

**Lucina Q. Uddin:** Data Curation, Methodology, Writing - Review & Editing

**Kuldeep Kumar:** Data Curation, Methodology, Writing - Review & Editing

**Gil Hoftman:** Data Curation, Methodology, Writing - Review & Editing

**Carrie E. Bearden:** Data Curation, Project Administration, Writing - Review & Editing, Supervision

Thank you to Ming Tsuang for contributions to the NAPLS2 project, Zailyn Tamayo for help accessing data, Ross D. Markello, Bratislav Misic et al. for the development of Neuromaps, and many thanks to all of the research participants who made this project possible.

## Data availability

Data are publicly available from the National Institute of Mental Health Data Archive: https://nda.nih.gov/edit_collection.html?id=2414 https://nda.nih.gov/edit_collection.html?id=2275

To facilitate reproducibility and rigor, analysis code is publicly available on GitHub: https://github.com/charles-schleifer/22q_chr_fmri

