## Supplemental Materials for "Unique functional neuroimaging signatures of genetic versus clinical high risk for psychosis"

**Supplemental Methods**

*Participants*

The neuroimaging dataset contains a total of 687 participants from two multi-site studies: 164 carriers of 22qDel along with 134 matched TD controls (Control-22q), and 240 individuals with CHR for psychosis plus 149 matched TD controls (Control-CHR).

22qDel and Control-22q data were shared from two scanners at UCLA (UCLAtrio and UCLAprisma), as well as the State University of New York in Syracuse, NY, USA, Sapienza University in Rome, Italy, and King's College London Institute of Psychiatry in London, UK. See Schleifer et al., 2023 for a full description of inclusion/exclusion criteria for 22qDel participants and matched controls (1). Briefly, a history of head injury or neurological disorder was exclusionary for all participants, and controls were excluded based on a personal history or first-degree family history of a psychosis spectrum illness. See **Supplementary Table S1** for participant counts and demographics by site.

CHR and Control-CHR data came from the North American Prodrome Longitudinal Study 2 (NAPLS2) (2,3), which includes eight sites in the United States and Canada. Study site details and inclusion criteria are described in Addington et al., 2012 (2). CHR status was defined by the Criteria of Psychosis-risk Syndromes based on the Structured Interview for Psychosis-Risk Syndromes (SIPS) (4). See **Supplementary Table S2** for participant counts and demographics by site.

| Site | Demographics | 22qDel | CONTROL | p |
|---|---|---|---|---|
| UCLAtrio | n | 51 | 38 | |
| | Age, Years, Mean (SD) | 17.1 (6.5) | 15.1 (4.2) | 0.106 |
| | Sex, Female, n (%) | 29 (56.9) | 19 (50.0) | 0.669 |
| UCLAprisma | n | 17 | 13 | |
| | Age, Years, Mean (SD) | 19.1 (8.9) | 19.9 (10.4) | 0.808 |
| | Sex, Female, n (%) | 9 (52.9) | 11 (84.6) | 0.152 |
| SUNY | n | 49 | 28 | |
| | Age, Years, Mean (SD) | 21.0 (2.3) | 20.8 (1.5) | 0.654 |
| | Sex, Female, n (%) | 21 (42.9) | 13 (46.4) | 0.948 |
| Rome | n | 19 | 21 | |
| | Age, Years, Mean (SD) | 27.4 (8.1) | 27.3 (6.4) | 0.970 |
| | Sex, Female, n (%) | 5 (26.3) | 14 (66.7) | 0.025 |
| KCL | n | 28 | 34 | |
| | Age, Years, Mean (SD) | 18.5 (6.6) | 17.7 (6.2) | 0.657 |
| | Sex, Female, n (%) | 13 (46.4) | 18 (52.9) | 0.799 |

**Supplementary Table S1.** Participant counts and demographics for all 22qDel sites. Each set of three rows contains the counts for each group and descriptions of the mean age and proportion of female participants for a given site, with p-values for comparisons between patient and control groups based on ANOVA or chi-squared tests, respectively.

| Site | Demographics | CHR | CONTROL-N | p |
|---|---|---|---|---|
| 01 | n | 37 | 18 | |
| | Age, Years, Mean (SD) | 18.9 (4.8) | 18.1 (3.6) | 0.518 |
| | Sex, Female, n (%) | 11 (29.7) | 7 (38.9) | 0.709 |
| 02 | n | 43 | 22 | |
| | Age, Years, Mean (SD) | 21.1 (5.1) | 21.1 (5.2) | 0.966 |
| | Sex, Female, n (%) | 22 (51.2) | 7 (31.8) | 0.222 |
| 03 | n | 24 | 23 | |
| | Age, Years, Mean (SD) | 18.7 (3.4) | 18.8 (4.5) | 0.949 |
| | Sex, Female, n (%) | 11 (45.8) | 11 (47.8) | 1.000 |
| 04 | n | 25 | 24 | |
| | Age, Years, Mean (SD) | 16.0 (1.7) | 17.1 (2.5) | 0.102 |
| | Sex, Female, n (%) | 12 (48.0) | 12 (50.0) | 1.000 |
| 05 | n | 47 | 25 | |
| | Age, Years, Mean (SD) | 19.5 (2.9) | 19.6 (2.9) | 0.875 |
| | Sex, Female, n (%) | 19 (40.4) | 12 (48.0) | 0.713 |
| 06 | n | 19 | 8 | |
| | Age, Years, Mean (SD) | 19.4 (4.3) | 21.8 (6.2) | 0.265 |
| | Sex, Female, n (%) | 6 (31.6) | 3 (37.5) | 1.000 |
| 07 | n | 39 | 14 | |
| | Age, Years, Mean (SD) | 16.7 (2.7) | 20.8 (5.3) | <0.001 |
| | Sex, Female, n (%) | 17 (43.6) | 8 (57.1) | 0.576 |
| 08 | n | 6 | 15 | |
| | Age, Years, Mean (SD) | 21.1 (5.2) | 21.6 (5.0) | 0.851 |
| | Sex, Female, n (%) | 1 (16.7) | 10 (66.7) | 0.112 |

**Supplementary Table S2.** Participant counts and demographics for all NAPLS sites. Each set of three rows contains the counts for each group and descriptions of the mean age and proportion of female participants for a given site, with p-values for comparisons between patient and control groups based on ANOVA or chi-squared tests, respectively.

*Neuroimaging acquisition*

UCLA 22q Prisma data were collected with Human Connectome Project (HCP)-style sequences on a Siemens Prisma 3 Tesla (3T) scanner. Four-hundred-and-twenty volumes (5.6 min) of resting BOLD data were acquired in 72 interleaved slices with multiband-8 acceleration (voxel size = 2 × 2 × 2 mm, TR = 800 ms, TE = 37 ms, flip angle = 52°, FOV = 208 × 208 mm), along with single-band reference images and a pair of spin-echo field maps with phase encoding in the anterior-posterior (AP) and posterior-anterior (PA) directions. T1w MP-RAGE and T2w SPC images were collected in 208 sagittal slices (voxel size = 0.8 × 0.8 × 0.8 mm, FOV = 256 × 256 mm) with (T1w TR = 2400 ms, TE = 2.22 ms) and (T2w TR = 3200 ms, TE = 563 ms).

UCLA and KCL 22q TimTrio resting BOLD data were acquired on a Siemens TimTrio 3T scanner in 34 interleaved axial slices (voxel size = 3 × 3 × 4 mm, TR = 2000 ms, TE = 30 ms, flip angle = 90°, FOV = 192 × 192 mm). Acquisition lasted 5.1 min and produced 152 volumes. High-resolution T1w MP-RAGE images were collected in 160 sagittal slices (voxel size = 1 × 1 × 1 mm, TR = 2300 ms, TE = 2.91 ms, flip angle = 90°, FOV = 240 × 256 mm)

SUNY 22q data were collected on a Siemens TimTrio 3T scanner. BOLD data were acquired in 34 axial slices (voxel size = 4 × 4 × 4 mm, TR = 2000 ms, TE = 30 ms, flip angle = 90°, FOV = 256 × 256 mm). Acquisition lasted 5.1 min and produced 152 volumes. High-resolution T1w MP-RAGE images were collected in 176 sagittal slices (voxel size = 1 × 1 × 1 mm, TR = 2530 ms, TE = 3.31 ms, flip angle = 7°, FOV = 256 × 256 mm).

22q data from Sapienza university in Rome were collected on a Siemens MAGNETOM Verio 3T scanner. (voxel size = 4 × 4 × 3 mm, TR = 3000 ms, TE = 30 ms, flip angle = 90°). 150 BOLD volumes were acquired. High-resolution T1w MP-RAGE (voxel size = 1 × 1 × 1 mm, TR = 2300 ms, TE = 2.98 ms, flip angle = 7°, FOV = 240 × 256 mm).

Details of BOLD acquisition for the NAPLS2 CHR study are described by Noble et al 2017 (3). Briefly, all data were acquired on comparable 3T scanners from either Siemens or General Electric. BOLD data were acquired with 30 4mm axial slices with a 1mm gap (TR = 2000 MS, TE = 30ms, flip angle = 77°, FOV = 220 x 220 mm). Acquisition lasted 5 min and produced 154 volumes. High-resolution 1 x 1 x 1.2 sagittal T1w images were also collected.

*Neuroimaging processing*

All data from 22qDel, CHR, and TD controls were processed with the same workflow, as described in detail in previous publications (1,5). Functional and structural images were processed with the Quantitative Neuroimaging Environment and Toolbox (6) to apply a modified version of the methods developed for the Human Connectome Project (HCP) (7) as well as motion scrubbing, i.e. censoring frames with displacement or intensity change thresholds exceeding those recommended by Power et al. (8–10). Functional connectivity analyses were computed on the residual of the signal after regression of motion time series, the mean signal time series from the ventricles and deep white matter, and the first derivatives of these measures.

*fMRI measures*

Three rs-fMRI measures were calculated for each scan: global brain connectivity (GBC), local connectivity (LC), and brain signal variability (BSV). All three measures used the same set of 360 cortical regions defined from multi-modal MRI in 210 healthy young adults from the HCP (11). Computations were performed in R using ciftiTools to manipulate neuroimaging data (12).

GBC is a well-validated measure defined as the average functional connectivity between a given brain region and all other regions (13,14). A high GBC value indicates a region in which signal is similar to many other regions of the brain, whereas a low GBC value represents a region that is dissimilar to the majority of other regions. GBC is sensitive to functional network disruptions in disorders such as schizophrenia (15,16). Here, we calculated this measure by computing functional connectivity (FC) between each region and each of the other 359 regions, followed by averaging the FC values for each region to achieve 360 unique GBC values (one per region). FC was calculated as the Fisher Z-transformed Pearson correlation between the mean BOLD time series in each region.

LC was calculated as a measure of the cohesiveness or homogeneity of vertex-level BOLD time series within each region. This approach is based on the network homogeneity method, wherein FC is computed between each pair of voxels in a chosen region (17). This is conceptually related to the regional homogeneity approach (18), except here homogeneity is calculated at the parcel level rather than for each voxel and its immediate neighbors. We generated a single average LC value for each of the 360 cortical regions by computing the full functional connectivity matrix between all vertices in a region and then taking the average of that.

BSV was calculated as the average temporal standard deviation of the BOLD time series in each region. This measure of variability is also referred to as resting state fluctuation amplitude.

To correct for variability related to site/scanner, we used neuroComBat (19), a neuroimaging-optimized implementation of the ComBat algorithm (20), which uses empirical Bayes methods to correct for batch/site effects with increased robustness compared to linear model approaches. Site correction was applied separately to the NAPLS2 data and the data from the multi-site 22qDel studies, and each of the three fMRI measures was corrected with a separate model. After ComBat, values for each measure were normalized within each region based on the mean and standard deviation for the relevant control group.

*Group-level fMRI comparisons*

For each fMRI measure, across each region, linear models were used to test the main effect of 22qDel versus matched controls (Control22q), and the main effect of CHR versus matched controls (ControlCHR). The models tested took the following form:

$$fMRI_{ij} \sim group_k + age + age^2 + sex + site + movement$$

In which the fMRI measure *i* (GBC, LC or BSV) at region *j* (1 of 360) is predicted by group *k* (either 22qDel versus Control22q, or CHR versus ControlCHR), controlling for linear and quadratic age, sex, site, and movement (measured as the percentage of frames scrubbed from each scan). p-values were computed for the main effect of group in each model, and were adjusted for False Discovery Rate (FDR) (21) across the 360 tests within each set of analyses for a given fMRI measure and reference group. Significance was evaluated at a threshold of FDR $q < 0.05$.

*Brain map comparisons*

In order to assess similarity of the three fMRI measures within and between clinical groups, we used permutation methods to compare the spatial brain maps for case-control comparisons with significant group main effects. For a given pair of maps, “map A” and “map B”, the comparison was conducted as follows: compute the Pearson correlation between the left hemispheres of maps A and B across 180 regions, repeat for the right hemisphere, and average to get the mean bilateral correlation. Next, use BrainSMASH to create 10,000 surrogate brain maps per hemisphere that preserve the original spatial autocorrelation structure of map A, and generate null models by testing the correlations between map B and each of these 10,000 surrogate maps per hemisphere. Two-tailed p-values were computed as the proportion of these 20,000 null-model values with an absolute value greater than the absolute value of the true average bilateral correlation between maps A and B.

We next used the same permutation testing procedure to compare the left hemisphere cortical maps from our case-control analyses to a set of 22 left hemisphere cortical maps from previously published datasets. Maps include metabolic and physiologic data from positron emission tomography (PET), gene expression from post-mortem tissue, and various measures from magnetoencephalography (MEG), structural MRI, and functional MRI; see **Table 2** for a description of each map (11,22–29). Analyses were restricted to the left hemisphere because some datasets did not include densely sampled right hemisphere data. All maps were transformed into the same left hemisphere surface-based parcellation.

For the majority of datasets, brain maps were processed with neuromaps, a toolbox containing resources for accessing, transforming, and comparing multi-modal brain datasets (30)**.** With this toolbox, multimodal published maps were transformed into the same coordinate space and re-parcellated based on the HCP multimodal atlas (11). For each threshold-free case-control fMRI left hemisphere map (e.g. the set of 180 coefficients for the effect of 22qDel versus

Control22q on BSV), correlations were tested with each of the 22 reference maps, and two-tailed p-values were computed from 10,000 BrainSMASH permutations and were evaluated at alpha = 0.05.

Spatial maps of gene expression from densely sampled microarray data from six typical adult post-mortem donors from the Allen Human Brain Atlas (31) were resampled to the same MRI atlas using the abagen toolbox (22). Spatial maps of typical parvalbumin (*PVALB*) and somatostatin (*SST*) gene expression across the left cortical hemisphere were generated as follows: Regional microarray expression data were obtained from 6 post-mortem brains (1 female, ages 24.0--57.0, 42.50 +/- 13.38) provided by the AHBA (32,33). Data were processed with the abagen toolbox (version 0.1.3; https://github.com/rmarkello/abagen) using a 360-region surface-based atlas in MNI space. First, microarray probes were reannotated using data provided by Arnatkevičiūtė et al. 2019 (34); probes not matched to a valid Entrez ID were discarded. Next, probes were filtered based on their expression intensity relative to background noise (35), such that probes with intensity less than the background in >=50.00% of samples across donors were discarded. When multiple probes indexed the expression of the same gene, we selected and used the probe with the most consistent pattern of regional variation across donors (i.e., differential stability) (32). The MNI coordinates of tissue samples were updated to those generated via non-linear registration using the Advanced Normalization Tools (ANTs; https://github.com/chrisfilo/alleninf). Samples were assigned to brain regions by minimizing the Euclidean distance between the MNI coordinates of each sample and the nearest surface vertex. Samples where the Euclidean distance to the nearest vertex was more than 2 standard deviations above the mean distance for all samples belonging to that donor were excluded. To reduce the potential for misassignment, sample-to-region matching was constrained by hemisphere and gross structural divisions (i.e., cortex, subcortex/brainstem, and cerebellum, such that e.g., a sample in the left cortex could only be assigned to an atlas parcel in the left cortex (34)). All tissue samples not assigned to a brain region in the provided atlas were discarded. Inter-subject variation was addressed by normalizing tissue sample expression values across genes using a robust sigmoid function (36). Normalized expression values were then rescaled to the unit interval. Gene expression values were then normalized across tissue samples using an identical procedure. Normalization was performed separately for samples in distinct structural classes (i.e., cortex, subcortex/brainstem, cerebellum). Samples assigned to the same brain region were averaged separately for each donor and then across donors, yielding a regional expression matrix. A similar procedure was used to produce the first principal component of gene expression published by Markello et al. 2021 (37) and accessed via neuromaps. A map of regional size was also generated for each of the 180 left hemisphere regions.

**Supplemental Results**

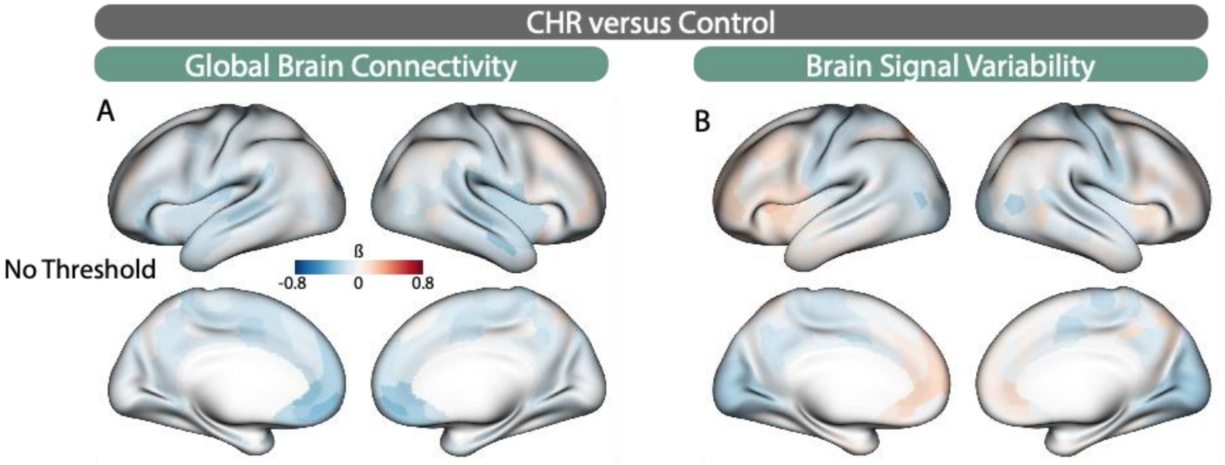


**Figure S1. Non-significant effects in CHR.** Threshold-free maps for group difference effects (CHR vs ControlCHR) for global brain connectivity and brain signal variability. No regions were significant at False Discovery Rate (FDR) q<0.05.

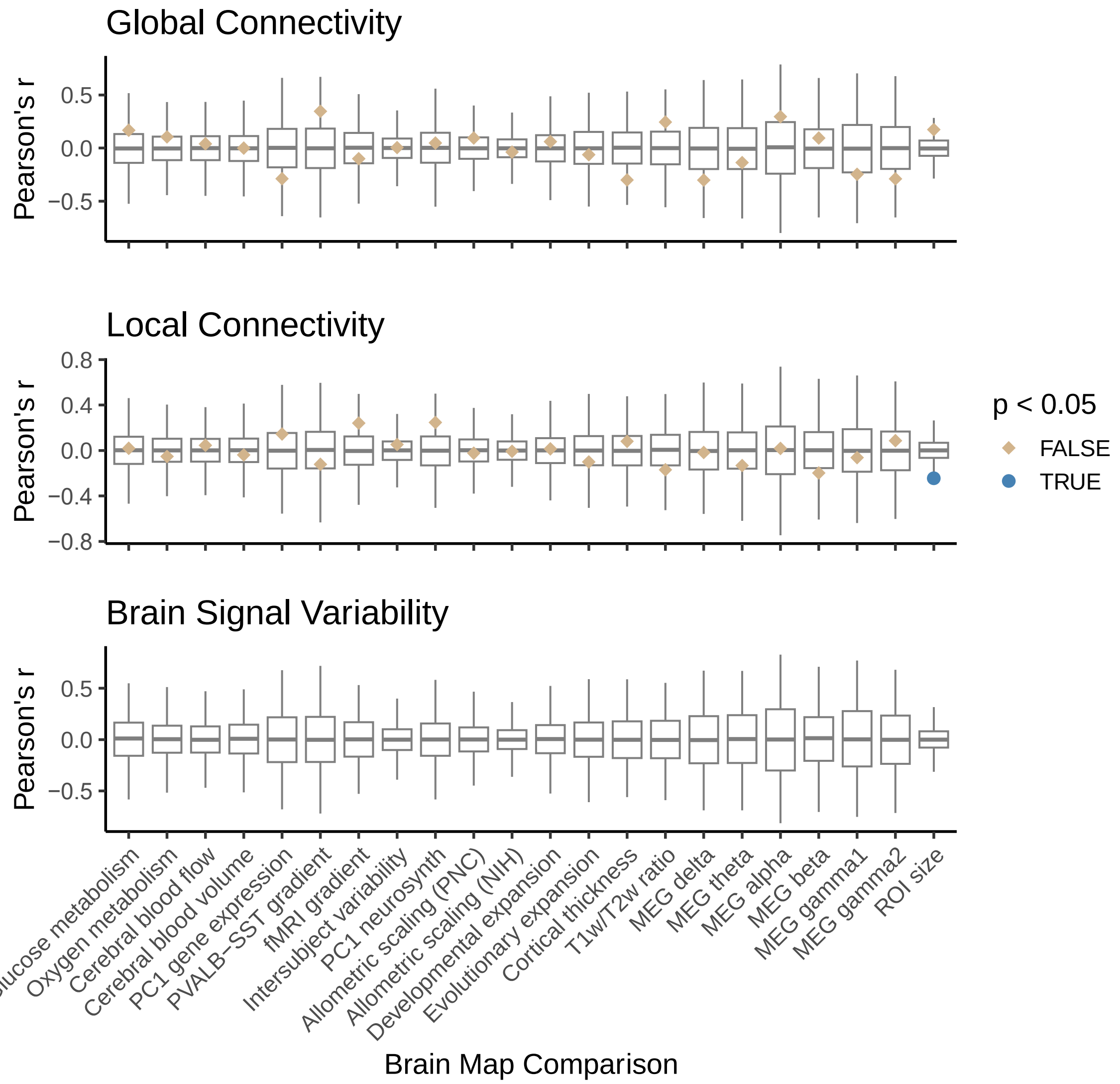


**Figure S2. Multi-modal brain map comparisons for CHR.** Left hemisphere cortical maps from CHR case-control models were tested for spatial similarity to multiple publicly available datasets including metabolic and physiologic data from positron emission tomography (PET), gene expression from post-mortem tissue, and various measures from magnetoencephalography (MEG), structural MRI, and functional MRI. Null distributions (gray box plots) were computed from the Pearson correlations between 10,000 spatial autocorrelation-preserving permutations of the target fMRI 22qDel-control map (named in the plot title) and the second map of interest (named on the x-axis). The true correlation values between the two maps of interest are marked by tan or blue points, and two-tailed p-values are computed from the proportion of values in the null distribution whose absolute value exceeds the absolute value of the true test statistic. Relationships are visualized for all three measures but should not be interpreted for global brain connectivity or brain signal variability which do not have sufficient effect sizes for CHR vs ControlCHR comparisons.

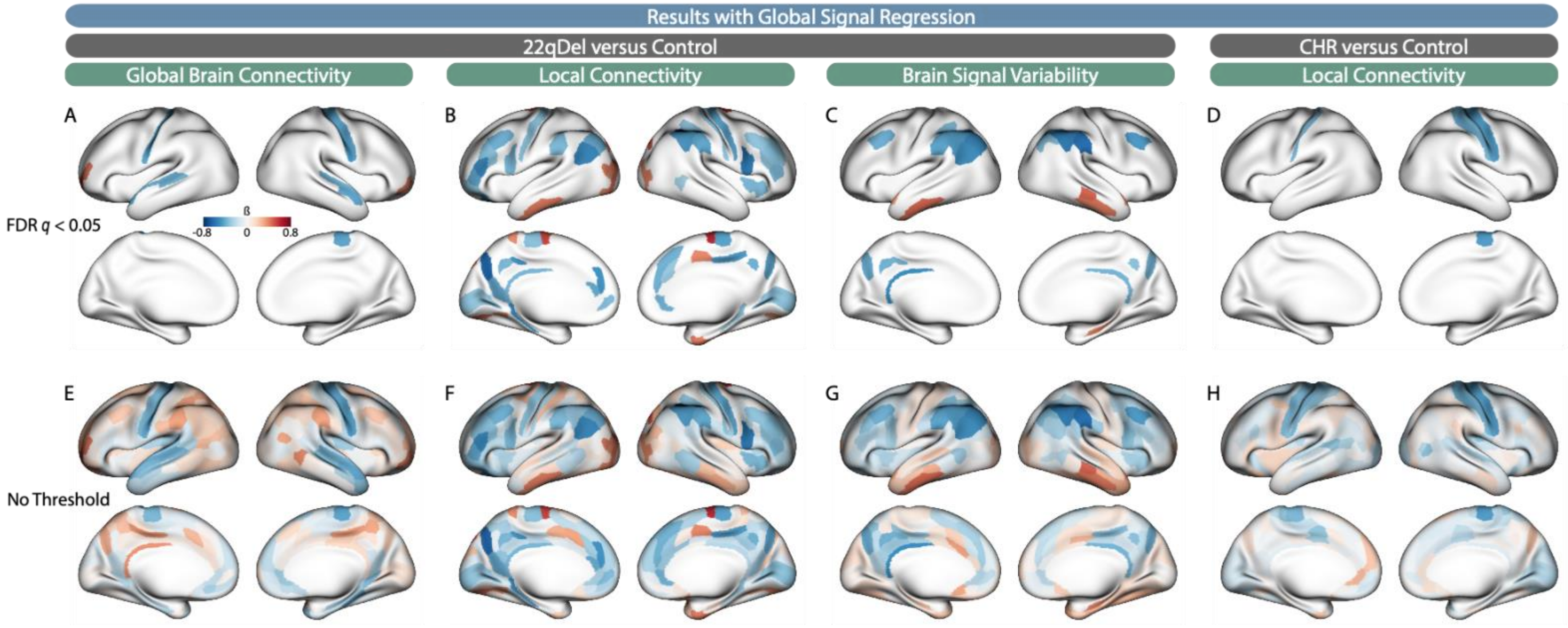


**Figure S3. Case-control fMRI results with Global Signal Regression (GSR).** Repeat of primary analyses using input fMRI time series that have been residualized based on the average whole-brain signal as well as the average signal from the ventricles and deep white matter, motion parameters, and their first derivatives. Results are highly consistent with our primary findings without GSR.
